# An Exploratory Multi-Session Study of Learning High-Dimensional Body-Machine Interfacing for Assistive Robot Control

**DOI:** 10.1101/2023.04.12.536624

**Authors:** Jongmin M. Lee, Temesgen Gebrekristos, Dalia De Santis, Mahdieh Nejati-Javaremi, Deepak Gopinath, Biraj Parikh, Ferdinando A. Mussa-Ivaldi, Brenna D. Argall

## Abstract

Individuals who suffer from severe paralysis often lose the capacity to perform fundamental body movements and everyday activities. Empowering these individuals with the ability to operate robotic arms, in high-dimensions, helps to maximize both functional utility and human agency. However, high-dimensional robot teleoperation currently lacks accessibility due to the challenge in capturing high-dimensional control signals from the human, especially in the face of motor impairments. Body-machine interfacing is a viable option that offers the necessary high-dimensional motion capture, and it moreover is noninvasive, affordable, and promotes movement and motor recovery. Nevertheless, to what extent body-machine interfacing is able to scale to high-dimensional robot control, and whether it is feasible for humans to learn, remains an open question. In this exploratory multi-session study, we demonstrate the feasibility of human learning to operate a body-machine interface to control a complex, assistive robotic arm in reaching and Activities of Daily Living tasks. Our results suggest the manner of control space mapping, from interface to robot, to play a critical role in the evolution of human learning.

## I. Introduction

People with cervical spinal cord injury (cSCI) can restore voluntary mobility using assistive robots. Assistive robots that are functional, intuitive, and learnable ensure continued robot use [1] and maximize opportunities for movement and independence to be restored [2]. While there are many strategies to support patients with robots, such as offloading tasks completely to full robot autonomy, technologies that empower patients to directly teleoperate complex robots offer the freedom to achieve everyday tasks with independence [3].

There are several examples of robotics platforms with the capability and control complexity to allow patients to perform Activities of Daily Living (ADLs) and Instrumental Activities of Daily Living (IADLs) via direct control [4]. However, the robotics community has yet to truly overcome the dimensionality mismatch problem that exists between interfaces and complex assistive robots. That is, there lies a mismatch between the number of control signal dimensions (1–3) that a single commercially-available interface is capable of issuing simultaneously, in comparison to the number of dimensions required for complex robot control (6 or more). Unfortunately, simultaneous and continuous control of all translation and orientation dimensions in complex robots has been extremely challenging with conventional interface solutions. This is critical because high control complexity is often needed in order for patients to achieve high-resolution dexterous movements in our physical world—and to efficiently perform everyday tasks, in a timely manner, with intention.

The problem of dimensionality mismatch and making high-resolution control accessible to patients can potentially be overcome using Body-Machine Interfaces (BoMIs). BoMIs use motion sensor technologies to measure movement from the surface of the body [5]. Unlike commercially-available interfaces, BoMIs have the capacity to generate control signal inputs, in high-dimensions, from residual body movements of even patients with cSCI (C3–C6) [6].

Moreover, BoMIs incentivize patients to use their remaining residual mobility [7]. For example, use of BoMIs have shown to increase patients’ upper body muscle strength and upper body mobility, with practice [6]. Thus, in addition to overcoming dimensionality mismatch, there are deeper implications that extend beyond assistance through robotics—that is, BoMIs also promote physical therapy and rehabilitation [8].

A current challenge with BoMIs is that we have a limited understanding of the extent to which people can learn to re-coordinate their body movements, to issue high-dimensional signals, necessary to control complex robots. Prior studies have raised concern that BoMI control could possibly be unintuitive and/or difficult to learn as control complexity in the robot increases [9]. In addition, the question of whether body movements can be executed with consistency and sufficient dexterity to complete functional tasks remains unanswered.

We take steps to address this challenge with an exploratory investigation of complex robotic arm operation via a high-dimensional BoMI. More specifically, in this paper, we make the following contributions:

- A presentation of a multi-session study of high-DoF robotic arm teleoperation via a high-dimensional BoMI.
- A demonstration that body machine interfacing is feasible and scalable to higher-DoF robots within a loosely-structured learning environment.
- An analysis of the impact of control space mappings on task performance, workload, and human learning.

The remainder of the paper is organized as follows. We summarize related literature in Section II. The methods of our multi-session study and its analysis are presented in Section III. The results of the study are reported in Section IV, with further discussion within Section V. In Section VI, we provide our conclusions.

## II. Related Work

Assistive machines range from low control complexity, such as mechanical wheelchairs, to high complexity, such as robotic arms. Robotic arms with a large number of degrees of freedom (DoFs) can be difficult to control with current commercially-available interfaces [10]. A common solution to this mismatch in dimensionality is modal control [11], in which only a subset of the control dimensions of the robot are operated at a given time. While modal control does facilitate access to the full control space, it does not allow people to access all control dimensions simultaneously and can lead to increases in time, cognitive load, and errors, when they attempt to perform tasks with complex robots [11].

People who suffer from upper and/or lower body paralysis experience loss of functional independence, and cascading effects further disrupt quality of life [2]. However, even in tetraplegia, some residual movements can remain intact [12]. Movement training and therapy that encourage residual movements and mobility are shown to promote neuroplastic changes and improve functionality [13].

BoMIs are able to capitalize on these residual body movements available to patients with paralysis. They are responsive to patient-to-patient variability (e.g., between or within levels of SCI) [5], enhance muscle strength and mobility [6], and achieve functional rehabilitation aims [8]. By casting a net of sensors on the body, the BoMI captures body movements. In general, BoMIs can be customized to individuals through their particular availability of body movements; for example, by tuning the BoMI map’s parameters (e.g., gains, offsets) session-by-session [6]. A classic approach to designing a BoMI map is to describe a linear relationship between sensor measurements and robot control commands [6]. More recent examples use iterative linear methods [14], feedback control (as opposed to feedforward control) [15], and deep learning methods such as adaptive nonlinear autoencoders [16].

Brain-Machine Interfaces (BMIs) using implants also offer the possibility of directly issuing high-dimensional control signals to overcome the problem of dimensionality mismatch. Although BMIs have enormous potential to help people with neurological disorders and injuries, including cSCI, they can be extremely invasive—requiring surgery to the brain and the permanent implantation of an electrode array [17]. Though BMI can be noninvasive using surface electroencephalography (EEG), there is less evidence that suggests the signal-to-noise ratio is sufficient for the continuous and simultaneous control of robots in high dimensions [18]. We focus in this work on the lesser-studied BoMI, because of its accessibility and demonstrated potential to promote motor recovery [6].

BoMIs are interfaced with a variety of assistive platforms, including powered wheelchairs [19] and robotic arms [20]— for all of which the control output is either discrete or the maximum continuous controllable dimensions are fewer than three. Using a BoMI for higher-dimensional and continuous control of assistive robots remains an open research question.

## III. Methods

Here we present the experimental details of our multi-session study of robotic arm operation using a high-dimensional BoMI (Figure 1).

**Fig. 1:**
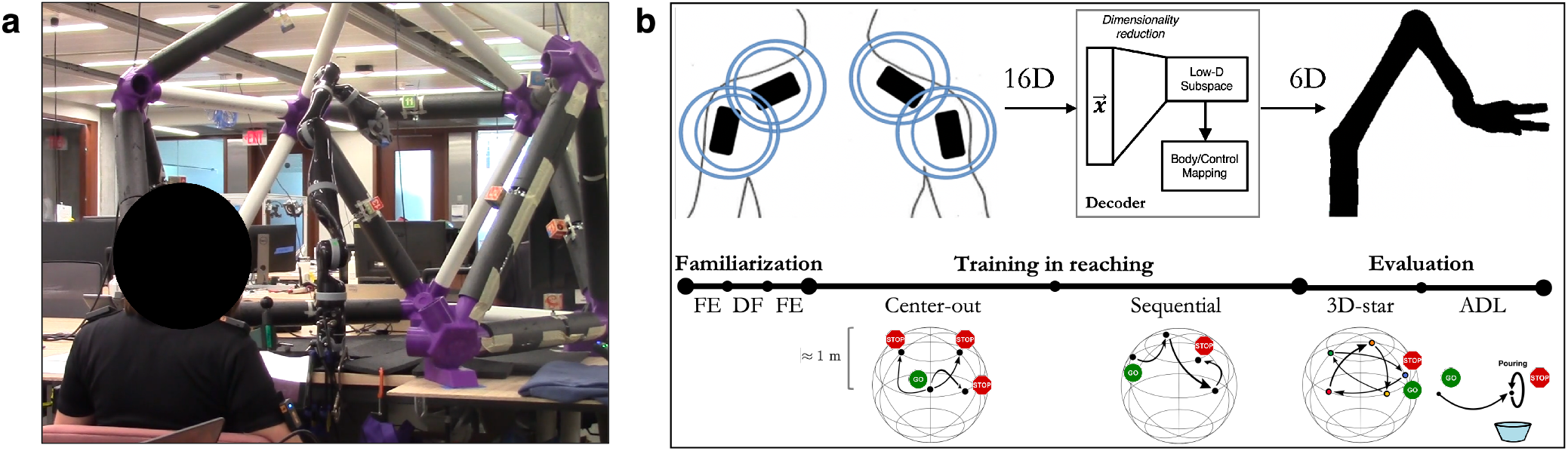
Overview of the interface-robot pipeline and study tasks. (a) Participant wearing the BoMI and operating the JACO robotic arm. Reaching targets affixed to a custom-built cage. (b) *Top*: BoMI IMUs (16D = 4 IMUs × 4D quaternion), mapped to a 6D linear subspace for continuous and simultaneous control of the robot in task-space (3D translation + 3D orientation) or joint-space (6D joint angles). *Bottom*: Progression of study session tasks: (1) free exploration (FE); (2) all-but-one DoF freezing (DF); (3) FE; (4) center-out reaching; (5) sequential reaching; (6) sequential reaching in a 3D-star shape; (7) ADL-inspired tasks.

### Participants

A total of ten adults (median age 28 ± 8 years) were recruited to participate in this study. Each participant completed five sessions of approximately two hours each, across five consecutive days. Participants were assigned to one of two groups: task-space (TS) or joint-space (JS). The TS group controlled the velocity of the robot end-effector in translation and orientation, while the JS group controlled the velocity of the robot joints. Group assignment was random and balanced. All sessions were conducted with the approval of the Northwestern University Institutional Review Board. All participants provided their written informed consent.

### Body-Machine Interface

A sensor net consisting of four inertial measurement unit (IMU) sensors (Yost Labs, Ohio, USA) are placed bilaterally on the scapulae and upper arms, and anchored to a custom shirt designed to minimize movement artifacts. Sensor placement is predetermined based on past BoMI studies. To maintain consistency between participants, we use orientation data from an additional reference chest sensor through a predetermined kinematic chain (chest → shoulders → upper arms). A Kalman filter is used as the filtering method for the IMUs, where orientation data is computed in real-time, onboard the IMU sensors, through a fusion of accelerometer and gyroscope measurements.

The pipeline and decoder design are visually represented in Figure 1b (top). The relative quaternion orientations of the four IMUs in the net (16D) are mapped (similar to [6]) to a 6D linear subspace using PCA. Initially, PCA is used to precompute a map (**A**) and an affine offset (**b**_0_), using data collected from an experienced user, performing a predefined set of upper body movements (shoulders forward/back and up/down; elbows in/out). The PCA map allows for a linear mapping between the 16D orientation data signal (**x**) and a 6D signal (**q**) as: **q** = **Ax** + **b**_0_.

### Robot Control

The lower-dimensional PCA subspace is used online to control a 7-DoF JACO robotic arm (Kinova Robotics, Quebec, Canada) with the fifth joint held fixed— which is the redundant joint in its kinematic chain. We hold this joint fixed so that both the TS and JS groups are operating the same number (6) of DoFs, and under the same control constraints.

The PCs of the lower-dimensional PCA subspace are mapped to the robot control space as follows. For the TS group, we prioritize control in translation over control in orientation by mapping the first three PCs (which by definition capture more body movement variance than the three latter PCs) to position (*x, y, z*) velocities, and the next three PCs to orientation (*θ* roll, *ϕ* pitch, *ψ* yaw) velocities. For the JS group, we map the PCs in the order of joints, in the kinematic chain of the robotic arm.

To avoid involuntary robot commands and to compensate for sensor noise during study trials, we use a control threshold formulation that is linearly proportional to the applied gains, shifted by a thresholding constant. To maximize the utility of the map for each individual, we customize the 6D signal (**q**) to the individual through scaling and shifting of control gains and offsets, determined through observation-based tuning. Control signals to the robot are published at an approximate rate of 100 Hz.

### Visual Feedback

A graphical-user interface (GUI) displayed on a tablet provides real-time visualization of the robot velocity commands to the participant. A scoring system also is displayed on the GUI to increase participant engagement and to provide trial-by-trial feedback on performance. Scores are calculated based on robot end-effector distance-to-target. Participants are told that this is a score but are not provided with the calculation details.

### Study Protocol

There are three phases of robot operation in the protocol of a single study session (Figure 1b, bottom).

1. **Familiarization**: The *free exploration* (FE) task encourages participants to explore and become familiar with the system. The *all-but-one DoF freezing* (DF) task iteratively introduces each control dimension, one at a time, while all other dimensions are kept frozen.
2. **Training**: The *center-out reaching* task starts always from a fixed center position when reaching targets.^1^ The *sequential reaching* task instead begins each reach from the prior reach’s target position. The order of targets is randomized and balanced across days to avoid ordering effects, and this order is preserved across participants.
3. **Evaluation**: To evaluate *sequential reaching*, participants are presented with five targets, that comprise a three-dimensional star, in fixed succession. To evaluate *ADL* task ability, participants: (a) transfer a cup (upside-down) from a dish rack and place it (upright) on the table, (b) pour cereal into a bowl, (c) scoop cereal from a bowl, and (d) throw away a surgical mask into a trash bin.

A trial ends upon successful completion or timeout. For a given reach to a target, success is defined within strict positional (1.00 cm) and rotational (0.02 rad, or 1.14°) thresholds, and the timeout is 90 seconds. For the ADL tasks, experimenters follow codified guidelines to determine when tasks are completed with a task timeout of 3 minutes. Over the course of the study, data from a total of 400 center-out, 400 sequential, and 250 3D-star reaching trials are gathered, as well as from 80 ADL task trials.

### Performance metrics

To evaluate the study’s reaching tasks and ADL tasks, we define the following performance metrics:

- Success rate

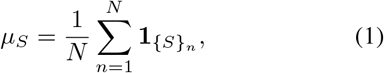

where

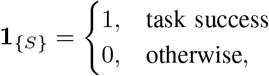

*N* is the total number of trials, and 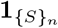 is an indicator function that summarizes task success.
- Successful completion time

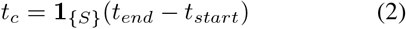
- Average number of collisions

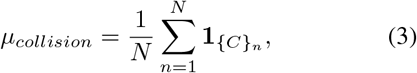

where 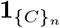 is an indicator function that summarizes task collisions.
- Normalized path length in a given reach

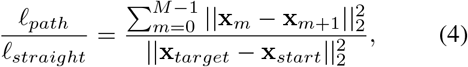

where *M* is the total number of samples (*f*_*s*_ = 10 Hz), **x**_*m*_ is the end-effector pose at the *m*^th^ sample, and **x**_*start*_ = **x**_0_.
- Average proportion of time spent within *k* percent of reach distance

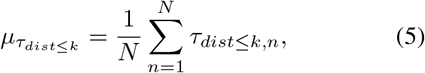

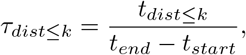

where *k* ∈ [0, 100]%. Note, we can substitute *dist* ≥ 100% to compute this beyond 100% of the reach distance (*τ*_*dist≥*100%_).

### Questionnaires

The NASA task load index (NASA-TLX) is an assessment tool to measure subjective workload in human-machine interfacing contexts [21], and is administered at the end of each session.

## IV. Results

We report results from the two evaluation tasks: sequential reaching (3D-star task) and activities of daily living (four ADL tasks). Our results find the manner of control space mapping (TS control versus JS control) to play a major role in both task performance and perceived workload.

### A. Task Performance

#### More Intuitive vs. More Learnable

The performance results from the ADL tasks are shown in Figure 2. Over five days, we observe that both the TS group and JS group do improve in success rate. The initial performance of the joint space group (JS) is superior to that of the task space group (TS)—specifically, on day 1, the JS group’s median success rate is higher (*p <* 0.05, Wilcoxon signed-rank test) and median trial time is lower. However, only TS demonstrates statistically significant improvements, between days 1 and 5, in success rate (*p <* 0.01) and trial time (*p <* 0.01). Thus, TS control appears to have a *greater capacity for improvement*, as measured by task success and trial time, while JS control demonstrates *greater success with naïve use*.

**Fig. 2:**
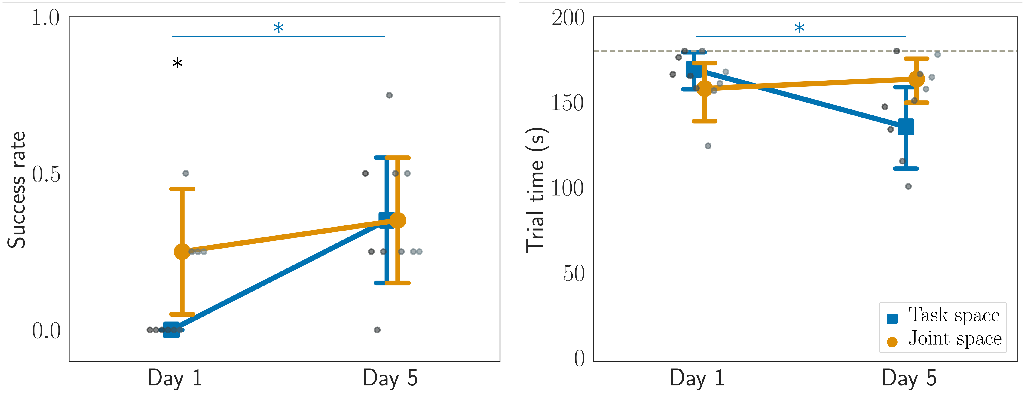
Success rate (left) and trial time (right) for ADL tasks on first and last days. Grayscale dots represent the mean value for each participant (TS: dark; JS: light), and the dotted line (right) represents a timeout of 180 seconds. The standard interquartile ranges are shown. ^*\**^*p <* 0.05.

Similarly, despite a noticeable early superiority in JS, on day 1, TS is able to reduce their number of collisions, compared to JS (days 1–4, *p <* 0.05; Kruskal-Wallis H-test), while collision numbers remain largely static for JS (Figure 3).

**Fig. 3:**
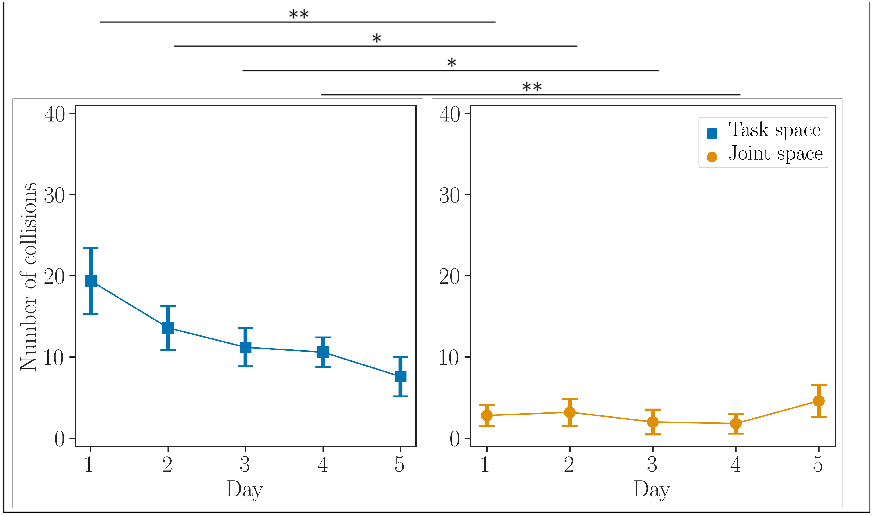
Average number of collisions during the 3D-star task over five days. Standard deviation is shown. ^*\**^*p <* 0.05, ^*\*\**^*p <* 0.01.

#### More (Room for) Improvement

Next, we evaluate success on the sequential reaching (3D-star) task. We examine how much time participants spend in workspace regions of interest: specifically, the proportion of time spent within 10% of the reach distance (*τ*_*dist≤*10%_) and beyond 100% of the reach distance (*τ*_*dist* ≥ 100%_ or farther from the target than is the starting position).^2^ The results are shared in Figure 4. Note that while for an ideal reach *τ* _*dist≥*100%_ would be zero and *τ*_*dist≤*10%_ minimized, *during learning* an *increase* in *τ*_*dist≤*10%_ appears to be a marker of improvement when targets are not yet achievable.

**Fig. 4:**
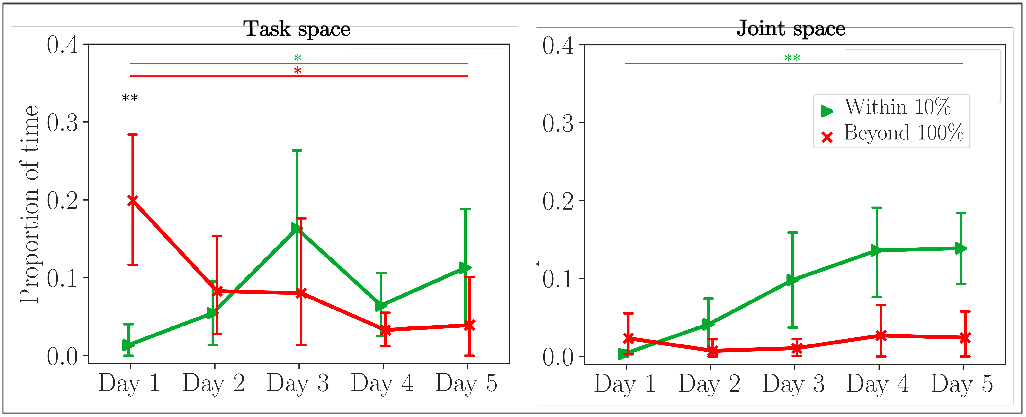
Proportion of time the robot end-effector spends within 10% of reach distance (green) and outside of 100% of reach distance (red) in the 3D-star task. The standard interquartile ranges are shown. ^*\**^*p <* 0.05, ^*\*\**^*p <* 0.01.

We also observe that, on day one, both groups spend more time beyond the starting distance (TS: 18 s; JS: 1.8 s) than near the targets (TS: 0.31 s; JS: 1.2 s). Both TS and JS groups significantly improve the amount of time spent near the targets (*τ*_*dist≤*10%_) between days 1 and 5 (*p <* 0.05; Kruskal-Wallis H-test). TS also significantly reduces how much time is spent beyond the starting distance (*τ*_*dist≥*100%_), between days 1 and 5 (*p <* 0.05), whereas this stays static in JS, largely due to having less room to improve.

### B. Perceived Workload

#### A Different Sort of Learning

Figure 5 shows the NASA-TLX assessment and the evolution of scores for subjective workload across sessions. We observe a marked reduction over sessions in perceived workload (median NASA-TLX score) for the JS group. By contrast, we observe only a slight decrease in perceived workload for the TS group. Furthermore, the perceived workload of the JS group is consistently lower than that of the TS group across days, and even on day one, when the JS group’s performance also was higher.

**Fig. 5:**
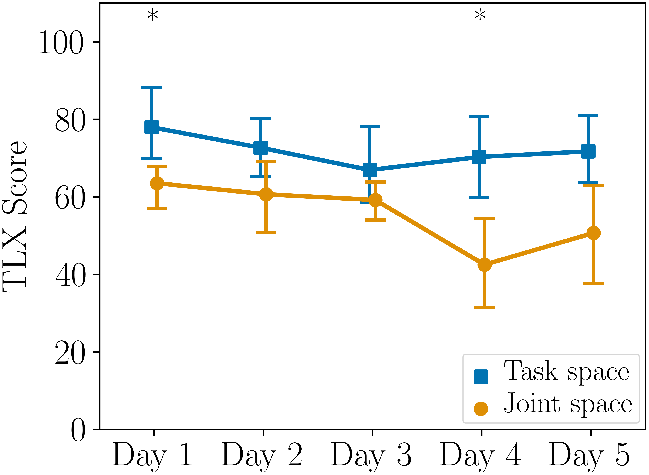
Comparison of subjective workload, measured via NASA-TLX, between study groups. Evolution of NASATLX scores over study sessions. The standard interquartile ranges are shown. ^*\**^*p <* 0.05.

To evaluate statistical significance between the two groups, we initially use the Kruskal-Wallis H-test to find a main effect, and Conover’s *post hoc* pairwise test, with Bonferroni adjustments, to make appropriate corrections. Only on days one and four do we find statistically significant differences between the two groups (*p <* 0.05).

We recall that the JS group did not improve much according to either of the ADL task performance metrics of success or trial time. While the JS group does not improve significantly in task performance, the group *does* improve in perceived workload—which perhaps is indicative of learning, albeit of a different sort than task performance learning (or at the very least familiarization).

We also perform a linear regression analysis between NASA-TLX scores and ADL task performance (success rate and trial time) and find that there are no strong correlations.

#### Learning Takes Work

The TS group does not improve measurably with respect to perceived workload. This group, however, does improve according to both performance metrics, of success and trial time. Thus, a possible explanation is that the gains in performance are expensive to acquire— simply put, learning takes work.

## V. Discussion

For individuals with paralysis, body-machine interfacing offers a promising path to increase agency, by enabling them to directly teleoperate high-dimensional assistive robots. To operate a BoMI requires learning a remapping of body movements to robot control signals. We have demonstrated this remapping within a 6D space to be learnable by an uninjured population, who were able to perform ADL tasks, in five days, under both control space mappings. Furthermore, this demonstration of 6D operation challenges the dimensionality mismatch problem that so often presents in the direct control of complex robotic arms, allowing for continuous and simultaneous operation of all robot control dimensions via the interface.

A focal question for our exploratory study was the feasibility of human learning to control high-dimensional robots, continuously and simultaneously, using a BoMI. Although several prior studies [6] had demonstrated the ability of people with severe paralysis to use a BoMI to complete 2D control tasks (without robot autonomy), other work had highlighted how challenging the control of additional dimensions can be, where 3D control took 2–4 times longer to learn than 1D or 2D control, in a simple virtual reaching task [9]. Therefore, the overall learning burden was expected to be nontrivial, especially given the novelty of the interfacing and the non-anthropomorphic robotic arm.

We furthermore have found that a multi-session study was necessary to tease out our learning results. That is, a one-session study would have shown evidence that JS control was superior to TS control in operating high-dimensional robots with a BoMI, and that TS control was unlearnable (Figures 2, 3, and 4). Instead, over five sessions, we have observed that TS control in fact was *more learnable* than JS control, which remained largely static over time. It moreover is likely that the TS group’s task performance would have continued to improve in a longer study and that we have not yet shown the full capability of human learning on this system. We expect that further gains might be possible with a more regimented learning curriculum that further customizes the presentation of tasks and control access to each individual. Where the inflection point exactly lies on the human learning curve in TS control remains a topic for further investigation.

Our analysis of the role of control space mappings in learning high-dimensional BoMI and robot control has provided evidence that *these mappings lead to different learning profiles* (Figures 2, 3, and 4). A possible explanation for why participants in the JS group initially intuit robot control more is the comparative simplicity of single-joint movements. A similar phenomenon is observed in patients with cerebellar ataxia, where lesions in the cerebellum cause patients to think out individual joint movements rather than being able to coordinate multi-joint movements [22]. In addition, TS is forced to also learn the forward kinematics of the robotic arm (whereas JS did not).

Not only do the learning profiles differ with respect to task proficiency, they also differ in regards to *what was learned*. We typically assess learning with respect to task performance. Equally important within the field of assistive and rehabilitation robotics, however, is the burden on the human operator. Learning to interface with the robot with a lower workload also is learning, and it achieves one of the driving motivators for the development of assistive robots. Broadly, this is critical when such robots have had a history of acceptance issues by their users [1].

Lastly, a significant limitation to this work is that our investigation considers only uninjured populations. Recall that the key motivation for this work was to obtain a baseline understanding of the feasibility, scalability, and learnability of high-dimensional robot teleoperation using a BoMI. While the BoMI previously had been shown to be effective at adjusting to the available residual movements in patients with cSCI (for 2D control) [6], questions related to the ability of human users to learn to teleoperate a robotic arm in high-dimensions using a BoMI, and whether control space mappings have any effect towards facilitating learning, were unanswered. Having now determined a baseline for human learning, our next steps will be to build on this work and apply the gained knowledge to a diverse population of patients with severe SCI, in longer studies.

## VI. Conclusions

Individuals with paralysis can benefit from assistive robotics and body-machine interfaces that help contribute to the return of their functionality and independence. In this paper, we share insights from a multi-session exploratory study that integrates body-machine interfacing with a high-dimensional assistive robot and investigates human learning to control this system. Our most significant finding was the impact of the control space mapping from interface to robot control. While joint-space control was found to be *more intuitive* prior to training, task-space control was subject to greater improvement over time and thus presented as *more learnable* with respect to task proficiency. That said, workload reduction is another critical aspect to human learning to interface with robots, and, in this regard, joint-space control exhibited superior learning capacity. Therefore, there appears to be a trade-off between intuitiveness and learnability when comparing the two control space mappings. Both of these learning curves merit further investigations—with patients with motor impairments and with longer studies—that more deeply probe their potential points of inflections and plateaus.

## ACKNOWLEDGMENT

Research reported in this publication was supported by the National Institute of Biomedical Imaging and Bioengineering (NIBIB), the NIH T32 Training Program and Eunice Kennedy Shriver National Institute Of Child Health & Human Development (NICHD), the National Science Foundation (NSF), National Institute on Disability, Independent Living and Rehabilitation Research (NIDILRR), and European Union’s Horizon 2020 Research and Innovation Program under the Marie Sklodowska-Curie, Project REBoT. The content is solely the responsibility of the authors and does not necessarily represent the official views of the National Institutes of Health.

## Footnotes

1 Ten targets are used as reaching goals, placed to maximize workspace coverage and diversify reaching movements. Placements remain fixed throughout the study and across participants.

2 A simple binary result of success is not informative, as no participants achieved the target location within our positional (1.00 cm) and rotational (0.02 rad or 1.14°) constraints on success. We also find the proportion of time metrics to be more informative than path length, for which no discernible trends emerge.

